# Cross-species connectome comparisons reveal the network attributes of memory capacity and time series prediction

**DOI:** 10.1101/2025.10.29.685101

**Authors:** Yazhe Yan, Geyang Wu, Haoming Yu, Qilin She, Yu Qian

**Author notes:** These authors contributed equally (Yazhe Yan, Yu Qian). These authors jointly supervised this work: Yazhe Yan, Yu Qian.

## Abstract

The brain’s connectome provides a powerful blueprint for designing efficient neural networks, yet the impact of incorporating its intricate, non-random architecture into machine learning models remains underexplored. Here, we integrate empirical structural connectomes from four model species—fruit fly, mouse, rat, and macaque—as the recurrent layer in echo state networks (ESNs). We demonstrate that biologically realized networks, particularly the macaque connectome, achieve superior performance in chaotic time series prediction and exhibit higher memory capacity compared to randomly shuffled controls. This computational advantage correlates with small-world topology, which scales with phylogenetic level. Crucially, we identify that weakly connected but highly central nodes are essential for optimal network dynamics; their targeted perturbation significantly degrades performance. Furthermore, functional connectomes from Alzheimer’s disease patients show computational deficits resembling those induced by weak-tie disruption in healthy networks. Our findings establish that evolved connectome topology is fundamental to efficient information processing, providing key principles for bio-inspired artificial intelligence.

## INTRODUCTION

The intricate network architecture of the brain, known as the connectome, has long served as a blueprint for understanding biological intelligence^1^. In contrast to artificial neural networks based on homogeneous architectures, biological brains exhibit heterogeneous connectivity patterns refined over millions of years of evolution^2^. These naturally evolved circuits demonstrate unparalleled efficiency in tasks that require complex parallel processing, adaptive learning, and energy-efficient computation^3^. Recent advances in connectome, particularly high-resolution mapping of neural circuits in model organisms such as *C. elegans*^4^, *Drosophila*^5^, and mouse^6^ have revealed design principles that challenge conventional machine learning paradigms. One promising approach to uncover the underlying mechanisms of this sparse, nonrandom, yet functionally optimized wiring is to relate structural connectivity to neural dynamics and the emergence of functional connectivity patterns^7^.

Building upon these insights, reservoir computing (RC) has emerged as a prominent paradigm for understanding how artificial recurrent neural networks extract information from continuous streams of external stimuli. Often implemented as echo state networks (ESNs) or liquid state machines, RC represents a promising platform for biologically inspired computation^8,9^. Using fixed recurrent connections, ESNs generate high-dimensional dynamical systems, thereby avoiding the intensive weight adjustment that characterizes conventional deep learning models^10^. Foundational studies have shown that even random recurrent connectivity can impart substantial computational power to neural networks^11,12^. More recent investigations suggest that integrating biological connectome data into reservoir architectures can further enhance performance in temporal processing and pattern recognition tasks^10^. For example, reservoirs based on the C. elegans connectome have been shown to outperform random networks in image classification tasks^4^, while reservoirs derived from the fruit fly connectome have produced superior results in predicting chaotic time series^13,14^. These empirical findings resonate with theoretical frameworks that posit that biological connectivity patterns inherently satisfy the separation and approximation properties essential for effective RC.

In addition, extending this line of inquiry, additional insights have emerged into the dynamic and computational capacities of neural systems. Quantitative evaluations of information processing in dynamical systems indicate that network structures similar to those in the brain can optimize both memory retention and nonlinear computation^15^. In parallel, network neuroscience has offered a comprehensive framework for understanding how structural connectivity underpins functional dynamics, leading to models that predict functional connectivity of the resting state from anatomical data^16^. Complementary studies in neuroimaging and computational modeling have also examined the effects of coupling, delay, and noise on brain dynamics^17,18^. Moreover, dynamic models of large-scale brain activity underscore the roles of criticality, multistability, and transient attractors in supporting complex information processes^19^. These models deepen our understanding of brain function and provide guiding principles for constructing artificial networks that mimic the adaptability of biological systems. In particular, characteristics such as the small world topology^20^, observed in biological networks, have been applied to ESNs to improve their robustness and learning performance^21^.

In this study, we present a novel approach that integrates the structural connectivity of biological brains into the ESNs reservoir layer. Drawing on previous work with connectome-based reservoirs and insights from cross-species connectomes, we have developed an optimization methodology that tailors the reservoir connection matrix according to biologically inspired principles. Our approach leverages the natural balance of segregation and integration observed in the brain, yielding significant improvements in temporal processing and dynamic adaptability compared to conventional ESN architectures. By addressing critical challenges, such as scaling microscopic connectomes to functional reservoirs and accounting for interspecies variability in network organization, this work establishes a robust framework for bioinspired reservoir computing that bridges neuroscience and artificial intelligence while outlining the potential for future neuromorphic applications.

### METHODOLOGY

#### Generating the Connectivity Matrix

We obtained connectivity matrices for fruit fly^22^, mouse^23^, rat^24^, and macaque^25^ from the Conn2Res toolbox ^26^, then scaled the initial biological matrices **W**_0_ by its spectral radius *ρ*(**W**_0_) to achieve optimal performance:

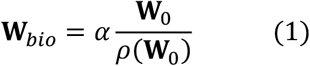

where *α* is a scaling constant that expresses *ρ*(**W**_0_), which impacts the reservoir’s echo state property. Standard ESNs possess the echo state property when their spectral radius satisfies *α* = *ρ*(**W**_*bio*_) < 1. For any species’ connectome, we can obtain the connectivity matrix **W**_*bio*_ after constraining the spectral radius, which will be used as the reservoir layer for subsequent performance tests.

#### The Echo State Network

A standard ESN typically consists of an input layer, a reservoir, and an output layer, as shown in Figure 1. The nodes and edges in Figure 1 represent neurons and synapses.

**Fig. 1.**
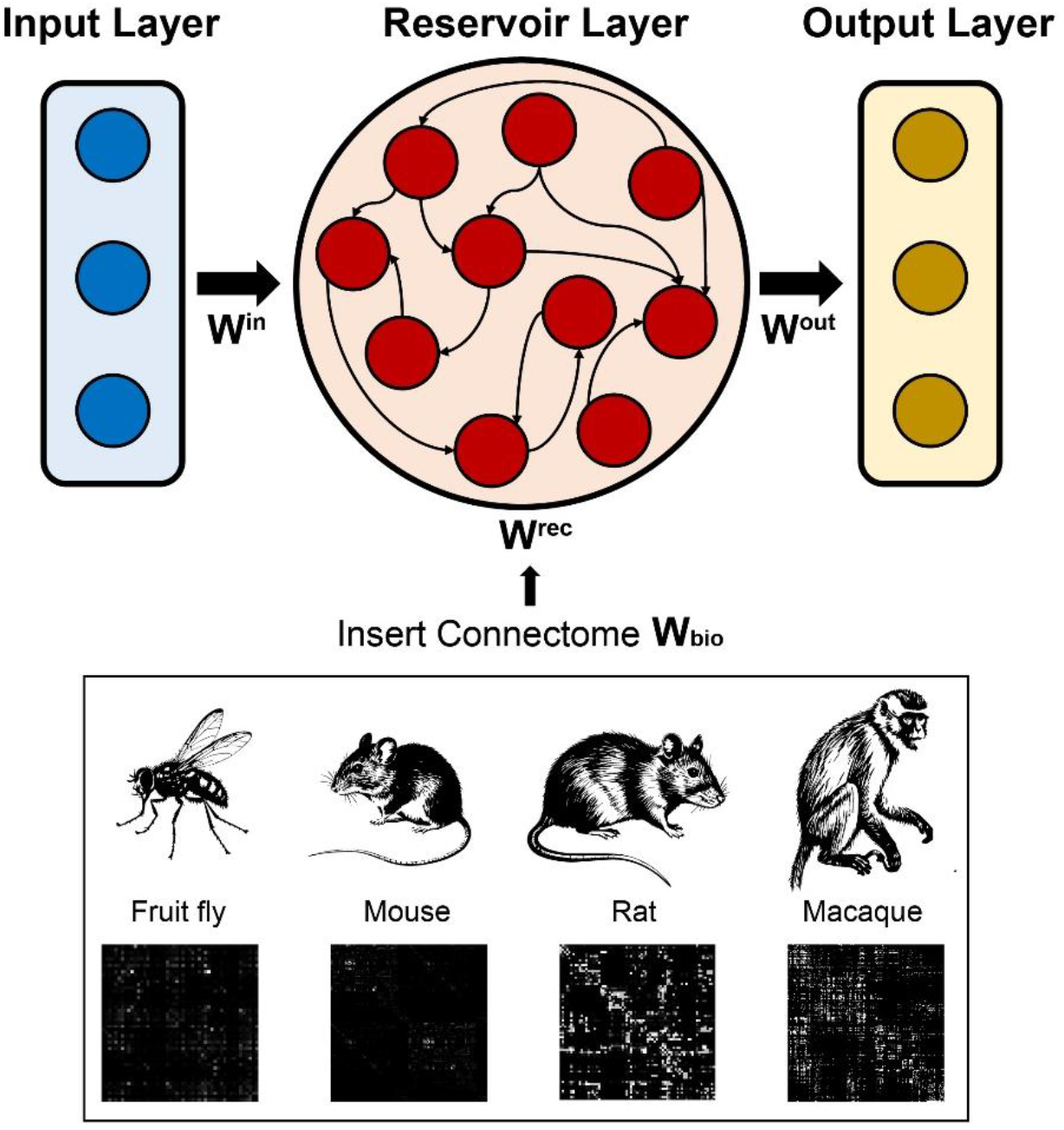
Employing animal connectomes as the recurrent layer in an Echo State Network. (Top) Schematic diagram of the ESN model, where the recurrent connectivity matrix (**W**^*rec*^) is replaced by the empirical brain connectomes of model organisms. (Bottom) The four model organisms used in this study (from left to right: fruit fly, mouse, rat, and macaque) and their corresponding structural connectome matrices.

The network dynamics of the ESN can be described as:

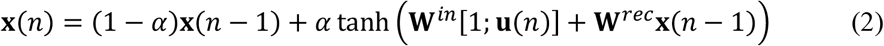

For some discrete time series input 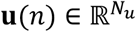, and known output 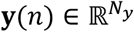 the ESN learns a prediction signal 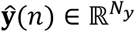, which minimizes *E*(**ŷ** (*n*), **y**(*n*)). Here [; ] denotes vector concatenation; **W**^*in*^ is the input weight matrix 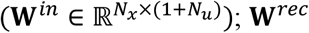 is the recurrent weight matrix 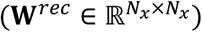, which is set according to **W**_*bio*_; **x**(*n* − 1) contains the previous reservoir activations 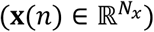; *α* is the leaking rate, (*α* ∈ (0,1]). The ESN is trained only at the readout layer. **W**^*out*^ is the output weight matrix 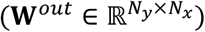. The output layer is defined as:

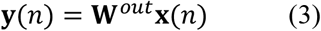

To solve this system, we use ridge regression (regression with L2-penalized regularization). Given the output of the known time series in matrix form **Y**, *λ* is the regularization coefficient, and **I** is the identity matrix:

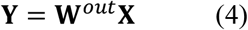

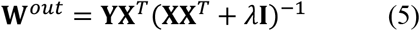

#### Time Series Prediction

We consider the Lorenz chaotic time series, which is used conventionally for benchmarking ESNs. The differential equation for the Lorenz time series is described below:

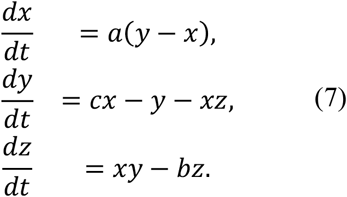

Here *a* = 10, *b* = 8/3, and *c* = 28 are used.

The objective of the time series prediction task for a series of size *T*, given *x*(*t*) for *t* ∈ [0,1, …, *m* − 1], is the correct prediction of *x*(*t*) for *t* ∈ [*m* + *r, m* + *r* + 1, …, *T*]. Here *r* controls how far into the future the model predicts; *m* ∈ ℤ^+^ and *r* ∈ ℤ^+^. We test all models to perform multi-step prediction on each time series and use a 5:80:15 warmup-train-validate split for all experiments.

#### Performance Evaluation

We evaluated the learning performance of ESNs in the tasks of memory capacity and prediction of non-linear time series. In the memory capacity task, an input series *u* is a one-dimensional random series sampled from a uniform distribution over [−1,1], and a desired series is the input series that is *r* time steps earlier. That is, the reservoir receiving a time series input *u*(*t*) at a time *t* was trained to output the predicted signal *y*(*t*) = *u*(*t* − *r*). This is why the reservoir has to store input information during the *r* time steps. For an output series **y** of a reservoir that learns an input series that is *r* steps earlier, the memory capacity (MC) is given as:

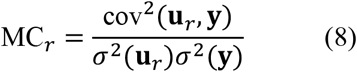

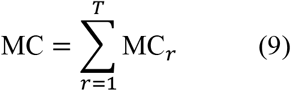

Finally, for a performance metric, we employed the Mackey-Glass and Lorenz chaotic time series for the reservoir performance evaluation and considered the Mean Squared Error (MSE) between each best-model prediction time series and the ground truth time series. The MSE is defined as the following:

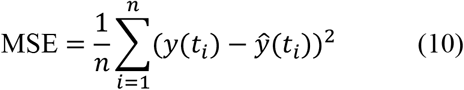

where *n* is the number of time steps; *y*(*t*_*i*_) is the labelled output at the time *t*_*i*_, and *ŷ* (*t*_*i*_) is the predicted output at time *t*_*i*_.

## Results

### Cross-species connectomes serve as high-performance reservoirs for temporal tasks

To investigate the computational principles of biological neural networks, we constructed echo state networks (ESNs) where the recurrent connectivity matrix was replaced by empirical structural connectomes from four model organisms: fruit fly, mouse, rat, and macaque (Fig. 1). We evaluated the performance of these bio-inspired ESNs on two key tasks: forecasting the chaotic Lorenz time series and memory capacity (MC).

Architectural differences between species’ connectomes conferred distinct computational advantages. The prediction accuracy, measured by mean squared error (MSE), and the memory capacity varied systematically with the scaling parameter α (Fig. 2a, b). For all species, a spectral radius around α = 1 yielded a favorable balance between nonlinear dynamics and stability, resulting in near-optimal performance for both tasks. A clear phylogenetic gradient in performance was observed: the macaque connectome consistently achieved the lowest MSE and the highest MC, followed by rat, mouse, and fly. This suggests that the evolutionary refinement of brain network topology enhances computational power for these temporal processing tasks.

**Fig. 2.**
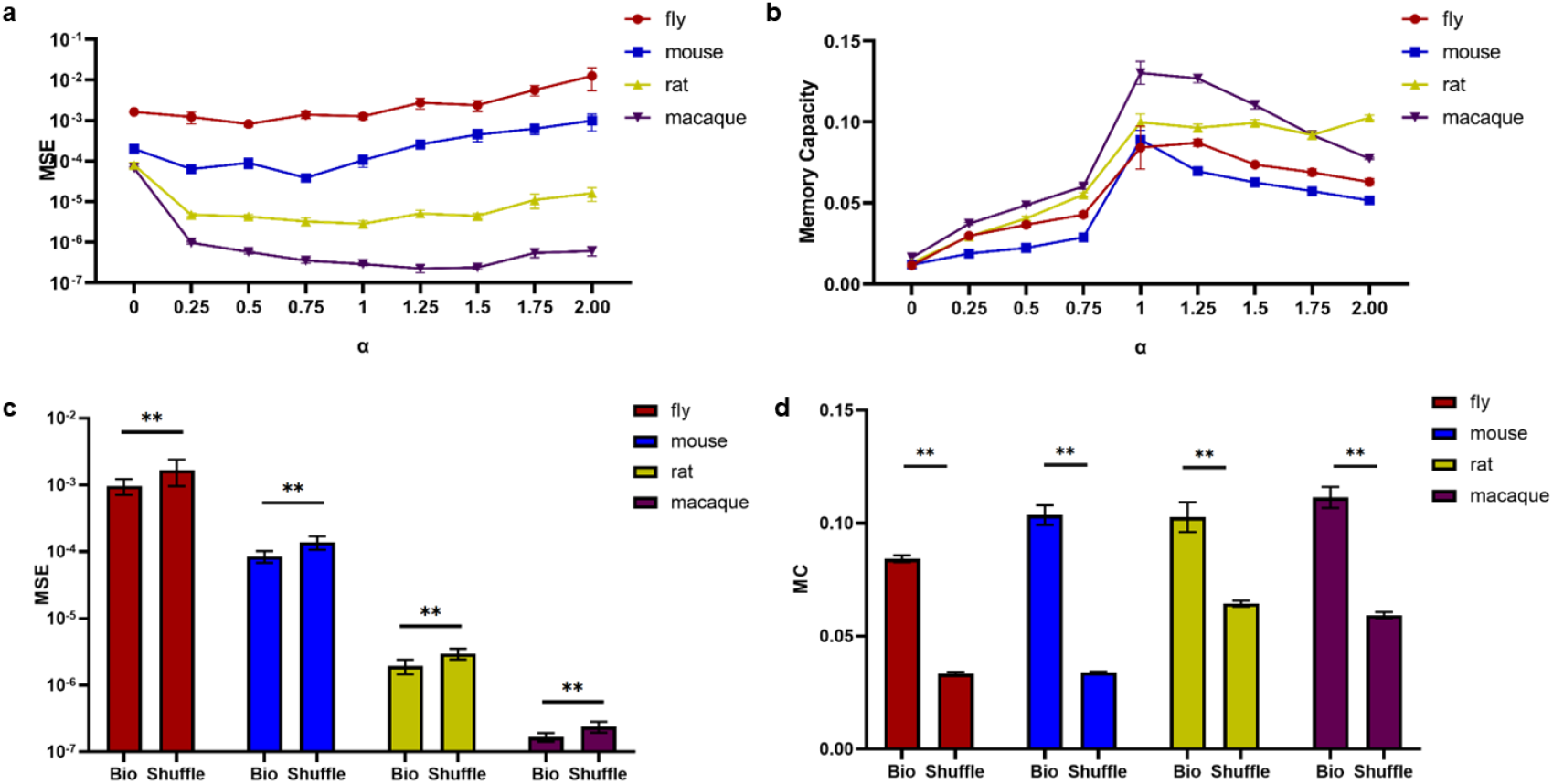
Computational performance of biological connectomes across four species in dynamical systems tasks. **a**, Mean squared error (MSE) of the Lorenz time series prediction task for fly, mouse, rat, and macaque connectomes, as a function of the scaling parameter α (range: 0-2). Lower MSE indicates better prediction performance. **b**, Memory capacity (MC) of the same connectomes as a function of α (range: 0-2). Higher MC indicates a greater ability to store and recall information. **c**, Comparison of Lorenz prediction MSE at α=1 between the original biological connectomes and their shuffled controls (where the connection matrix is randomized). Biological networks show significantly lower MSE than shuffled controls (paired t-test, p < 0.01 for all species). Error bars represent s.e.m. **d**, Comparison of memory capacity at α=1 between the original biological connectomes and their shuffled controls. Biological networks demonstrate a significantly higher memory capacity than shuffled controls (paired t-test, p < 0.01 for all species). Error bars represent s.e.m. *In c and d, *p < 0.05, **p < 0.01, **p < 0.001.

To ascertain that this performance was due to the specific biological wiring rather than generic network properties, we compared the native connectomes to shuffled controls, in which the connection weights were randomized while preserving the overall weight distribution. This shuffling procedure dramatically degraded performance across all species (Fig. 2c, d). The biological networks showed significantly lower MSE and higher MC than their shuffled counterparts (paired t-test, p < 0.01 for all species), underscoring that the non-random, evolved topology of the connectome is fundamental to its computational efficacy.

### Computational superiority correlates with small-world topology and depends on weak-tie nodes

Given the performance differences between species and the detrimental effect of shuffling, we sought to identify the specific topological features underlying efficient computation. We first computed the small-worldness (SW) of each network. The native connectomes exhibited a pronounced small-world structure, with SW increasing along the phylogenetic scale from fly to macaque (Fig. 3a). In contrast, shuffled controls showed consistently low SW (∼0.2), confirming that randomization disrupts this biologically prevalent architecture.

**Fig. 3.**
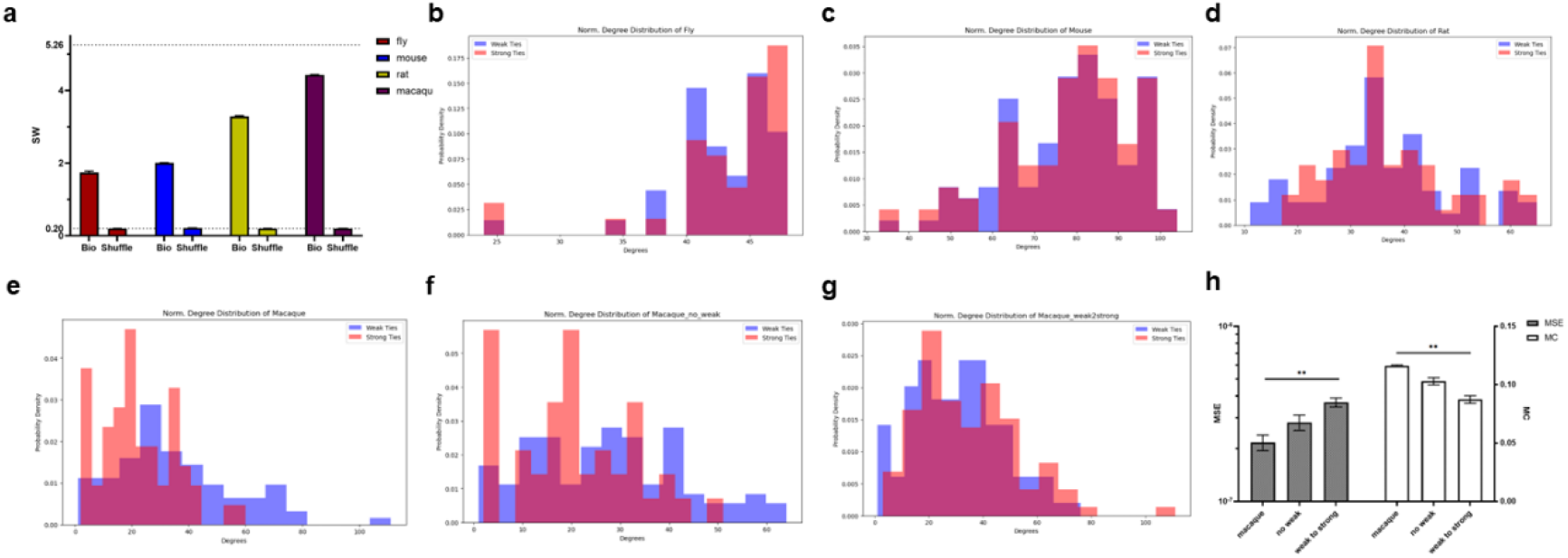
Small-world topology and the critical role of weak ties in biological connectomes. **a**, Small-worldness (SW) of native and shuffled connectomes across four species. SW was calculated as the ratio of normalized clustering coefficient to normalized characteristic path length (SW = (C/C_0_)/(L/L_0_)), where C and L are the clustering coefficient and path length of the network, and C_0_ and L_0_ are those of a random network. The native connectomes exhibit increasing SW with phylogenetic scale, a hallmark of small-world topology, whereas shuffled controls show consistently low SW (∼0.2). **b-e**, Degree distributions of strong-tie (red) and weak-tie (blue) nodes for the (b) fly, (c) mouse, (d) rat, and (e) macaque connectomes. Strong-tie and weak-tie nodes were defined as the top and bottom 10% of nodes based on their average connection strength, respectively. **f, g**, Altered degree distributions following network manipulations. f, Degree distribution after the removal of weak-tie nodes. g, Degree distribution after converting weak ties to strong ties. **h**, Impact of weak-tie manipulations on computational performance. A two-way ANOVA revealed that both network perturbations (removing weak ties and converting them to strong ties) led to a significant increase in MSE for the Lorenz prediction task and a significant decrease in memory capacity, underscoring the critical importance of weak-tie nodes for network performance. Data are presented as mean ± s.e.m. (n=10 networks per species). *p < 0.05, **p < 0.01, ***p < 0.001.

We hypothesized that nodes with weak connections, which are abundant in small-world networks, might play a disproportionately important role. We defined “strong-tie” nodes as those belonging to the top 10% of connection weights and “weak-tie” nodes as those in the bottom 10%. Analysis of the degree distributions revealed that strong-tie and weak-tie nodes are topologically distinct, particularly in higher-order species (Fig. 3b-e). In the macaque connectome, a substantial number of weak-tie nodes possessed high degrees, indicating that these are highly central, yet weakly connected, hubs within the network (Fig. 3e).

To functionally test the importance of these weak ties, we performed two network manipulation experiments: removal of the weak-tie nodes, and augmentation of their connection strengths to match the strong-tie range. Both perturbations severely altered the network’s degree distribution (Fig. 3f, g) and led to a significant performance deficit, manifesting as increased MSE and decreased MC (Two-way ANOVA, p < 0.001; Fig. 3h). This demonstrates that the native configuration of weak-tie nodes is critical for the computational capabilities of biological connectomes.

### Alzheimer’s disease impairs network performance by disrupting critical connectivity

Finally, we explored the translational relevance of our findings by examining functional connectomes derived from human neuroimaging data^27^. We obtained and analyzed pre-processed functional connectivity matrices from Cognitively Normal (CN) participants and patients with Alzheimer’s Disease (AD) (Fig. 4a). When deployed as reservoirs in our ESN framework, the AD connectomes exhibited significantly worse performance than the CN connectomes, with higher MSE for prediction and lower MC (t-test, p < 0.01; Fig. 4b).

**Fig. 4.**
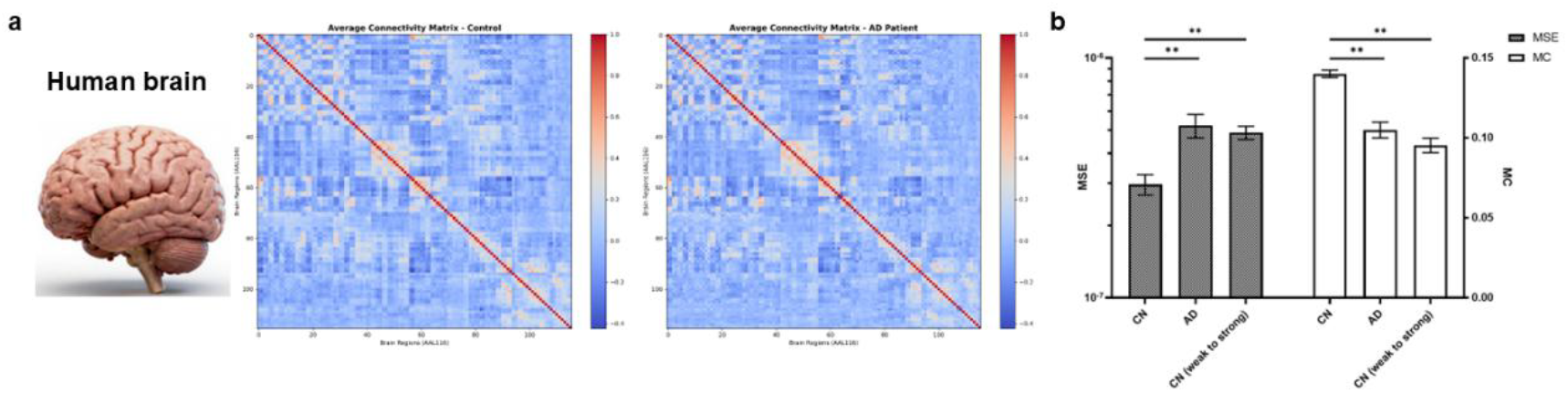
Degraded computational performance in functional connectomes of Alzheimer’s disease patients. **a**, Group-averaged functional connectivity matrices for Cognitively Normal (CN, left) and Alzheimer’s Disease (AD, right) participants. The AD matrix exhibits altered connectivity patterns compared to the CN group. **b**, Comparative performance on the Lorenz prediction (MSE, lower is better) and memory capacity (MC, higher is better) tasks. Performance is shown for three conditions: the native CN connectomes, the native AD connectomes, and the manipulated CN connectomes where weak ties were converted to strong ties. The AD connectomes show significantly impaired performance (higher MSE and lower MC) compared to the native CN connectomes (**p < 0.01, ***p < 0.001, t-test). Notably, the computational impairment in AD connectomes resembles the performance degradation observed in CN connectomes after the perturbation of weak ties. Data are presented as mean ± s.e.m.

Strikingly, the computational deficit in AD connectomes phenocopied the effect of perturbing weak ties in healthy networks. Artificially converting weak ties to strong ties in the CN connectomes caused a significant performance drop, bringing their performance closer to the impaired level observed in AD (Fig. 4b). This suggests that the pathophysiology of AD may involve a disruption of the delicate balance between strong and weak connections, compromising the network’s information processing capacity that relies critically on weak-tie architecture.

## Discussion

In Our study demonstrates that the naturally evolved architecture of the brain is computationally non-random. By integrating structural connectomes from four species into a reservoir computing framework, we found that their specific topological attributes—particularly small-world organization and weak-tie architecture—are crucial for efficient information processing in temporal tasks.

We show a clear phylogenetic gradient in computational performance, with the macaque connectome achieving superior results in both chaotic time series prediction and memory capacity (Fig. 2a, b). This aligns with the view that evolution shapes brain dynamics towards more efficient topologies. The significant performance degradation upon randomizing the biological connectomes (Fig. 2c, d) confirms that their advantage stems from their evolved fine-scale structure, not basic network statistics.

We attribute this advantage largely to small-world attributes, which systematically increased with phylogenetic scale (Fig. 3a). More importantly, we pinpoint a critical, previously underappreciated role for weakly connected nodes. In higher-order species like the macaque, many highly central nodes interact via weak ties (Fig. 3e). Our perturbation experiments revealed that these nodes are indispensable; disrupting them severely impaired network performance (Fig. 3h). We postulate that weak connections act as dynamic stabilizers or information bridges, facilitating flexible communication between modules without causing excessive synchronization, thereby enriching the reservoir’s computational repertoire.

Translating these principles to human neuroimaging, we found that functional connectomes from Alzheimer’s disease (AD) patients exhibit computational deficits resembling those induced by weak-tie disruption in healthy networks (Fig. 4b). This offers a novel network dynamics perspective on cognitive dysfunction in AD, suggesting its pathophysiology may partly stem from a breakdown of the critical weak-tie architecture essential for temporal information processing.

Some limitations of our study should be noted. The structural connectomes used vary in spatial scale and resolution, and our model is static, lacking dynamic mechanisms like synaptic plasticity. Future work incorporating spiking neuron models and plasticity rules could further unlock the potential of bio-inspired computing. Furthermore, the functional significance of “weak ties” warrants a more refined theoretical framework.

In summary, our work elucidates a clear path from identifying efficient biological network topologies through cross-species comparison, to dissecting their key structural features, and validating their functional importance in a human disease model. This research provides concrete design principles for building more adaptive, robust, and energy-efficient, bio-inspired artificial intelligence systems.

## Acknowledgments

We thank Chao-Ming Wang for suggestions and support. We appreciate the technical assistance from H. Zhang and Q. Hu. We acknowledge the assistance with the brain connectome processing from Z.X. Xia.

## Author contributions

Conceptualization: Y.Q., and Y.Y.; Code writing and brain simulation: Y.Y.; Methodology: Y.Y.; Investigation: Y.Q., Y.Y., and Q.S.; Data acquisition: Q.S., G.W., and H.Y.; Project administration: Y.Q., Y.Y., and G.W.; Manuscript writing: Y.H., Y.Y., and G.W. with inputs from others.

## References

1. Petersen, S. E. & Sporns, O. Brain Networks and Cognitive Architectures. Neuron 88, 207–219 (2015).

2. Pang, J. C., Rilling, J. K., Roberts, J. A., Van Den Heuvel, M. P. & Cocchi, L. Evolutionary shaping of human brain dynamics. eLife 11, e80627 (2022).

3. Laughlin, S. B. & Sejnowski, T. J. Communication in Neuronal Networks. Science 301, 1870–1874 (2003).

4. Bardozzo, F., Terlizzi, A., Simoncini, C., Lió, P. & Tagliaferri, R. Elegans-AI: How the connectome of a living organism could model artificial neural networks. Neurocomputing 584, 127598 (2024).

5. Morra, J. & Daley, M. Using Connectome Features to Constrain Echo State Networks. in 2023 International Joint Conference on Neural Networks (IJCNN) 1–8 (IEEE, Gold Coast, Australia, 2023). doi:10.1109/IJCNN54540.2023.10191832.

6. Chen, G., Scherr, F. & Maass, W. A data-based large-scale model for primary visual cortex enables brain-like robust and versatile visual processing. Sci. Adv. 8, eabq7592 (2022).

7. Suárez, L. E. Learning function from structure in neuromorphic networks. 3, (2021).

8. Maass, W., Natschläger, T. & Markram, H. Real-Time Computing Without Stable States: A New Framework for Neural Computation Based on Perturbations. Neural Comput. 14, 2531–2560 (2002).

9. Jaeger, H. The “echo state” approach to analysing and training recurrent neural networks – with an Erratum note.

10. Bianchi, F. M., Scardapane, S., Lokse, S. & Jenssen, R. Reservoir Computing Approaches for Representation and Classification of Multivariate Time Series. IEEE Trans. Neural Netw. Learn. Syst. 32, 2169–2179 (2021).

11. Verstraeten, D., Schrauwen, B., D’Haene, M. & Stroobandt, D. An experimental unification of reservoir computing methods. Neural Netw. 20, 391–403 (2007).

12. Dambre, J., Verstraeten, D., Schrauwen, B. & Massar, S. Information Processing Capacity of Dynamical Systems. Sci. Rep. 2, 514 (2012).

13. Tanaka, G. et al. Recent advances in physical reservoir computing: A review. Neural Netw. 115, 100– 123 (2019).

14. Morra, J. & Daley, M. Imposing Connectome-Derived Topology on an Echo State Network. in 2022 International Joint Conference on Neural Networks (IJCNN) 1–6 (IEEE, Padua, Italy, 2022). doi:10.1109/IJCNN55064.2022.9892629.

15. Sussillo, D. & Abbott, L. F. Generating Coherent Patterns of Activity from Chaotic Neural Networks. Neuron 63, 544–557 (2009).

16. Bassett, D. S. & Sporns, O. Network neuroscience. Nat. Neurosci. 20, 353–364 (2017).

17. Friston, K. Beyond Phrenology: What Can Neuroimaging Tell Us About Distributed Circuitry? Annu. Rev. Neurosci. 25, 221–250 (2002).

18. Deco, G., Kringelbach, M. L., Jirsa, V. K. & Ritter, P. The dynamics of resting fluctuations in the brain: metastability and its dynamical cortical core. Sci. Rep. 7, 3095 (2017).

19. Breakspear, M. Dynamic models of large-scale brain activity. Nat. Neurosci. 20, 340–352 (2017).

20. Watts, D. J. & Strogatz, S. H. Collective dynamics of ‘small-world’ networks. (1998).

21. Kawai, Y., Park, J. & Asada, M. A small-world topology enhances the echo state property and signal propagation in reservoir computing. Neural Netw. 112, 15–23 (2019).

22. Chiang, A.-S. et al. Three-Dimensional Reconstruction of Brain-wide Wiring Networks in Drosophila at Single-Cell Resolution. Curr. Biol. 21, 1–11 (2011).

23. Rubinov, M., Ypma, R. J. F., Watson, C. & Bullmore, E. T. Wiring cost and topological participation of the mouse brain connectome. Proc. Natl. Acad. Sci. 112, 10032–10037 (2015).

24. Bota, M., Sporns, O. & Swanson, L. W. Architecture of the cerebral cortical association connectome underlying cognition. Proc. Natl. Acad. Sci. 112, (2015).

25. Modha, D. S. & Singh, R. Network architecture of the long-distance pathways in the macaque brain. Proc. Natl. Acad. Sci. 107, 13485–13490 (2010).

26. Suárez, L. E. et al. Connectome-based reservoir computing with the conn2res toolbox. Nat. Commun. 15, 656 (2024).

27. Xu, J. et al. A Collection of Brain Network Datasets. 738521608128 Bytes The University of Auckland 10.17608/K6.AUCKLAND.21397377 (2024).

